# TgGloL is an atypical glyoxalase/VOC domain-containing apicoplast protein that is important for the growth of *Toxoplasma*

**DOI:** 10.1101/2024.09.04.611175

**Authors:** Syrian G. Sanchez, Esther Pouzet, Loïc Guimbaud, Arnault Graindorge, Laurence Berry, Sébastien Besteiro

## Abstract

Glycolysis is a conserved metabolic pathway that converts glucose into pyruvate in the cytosol, producing ATP and NADH. In *Toxoplasma gondii* and several other apicomplexan parasites, some glycolytic enzymes have isoforms located in their plastid (called the apicoplast). In this organelle, glycolytic intermediates like glyceraldehyde 3-phosphate (GAP) and dihydroxyacetone phosphate (DHAP) are imported from the cytosol and further metabolized, providing ATP, reducing power, and precursors for anabolic pathways such as isoprenoid synthesis. However, GAP and DHAP can spontaneously convert into methylglyoxal, a toxic by-product detoxified by the glyoxalase system, typically involving Glyoxalase-1 (Glo-1) and Glyoxalase-2 (Glo-2). In T. gondii, we identified an atypical protein, TgGloL, containing a Glo-1-like motif but with limited homology to typical Glo enzymes. TgGloL localizes to the apicoplast, and its conditional knockdown impairs parasite growth, indicating its importance. While a specific and direct role for TgGloL in methylglyoxal detoxification within the apicoplast remains unclear, it is crucial for maintaining organelle homeostasis and for overall parasite fitness.

## Introduction

Many parasites of the phylum Apicomplexa harbor a plastid that originated from a double endosymbiotic event. This organelle, called the apicoplast, hosts several important metabolic pathways that contribute to the fitness of the parasites (Kloehn *et al*., 2021). Some apicomplexan parasites, including *Plasmodium* and *Toxoplasma*, have a serious impact on human health. For instance, several *Plasmodium* species are causing malaria, a devastating disease responsible for more than half a million deaths each year, mostly in Africa (Ashley *et al*., 2018). On the other hand, infection with *Toxoplasma gondii*, the causative agent of toxoplasmosis, is only potentially fatal in immunocompromised individuals and developing fetuses, but the parasite is distributed worldwide and infects up to one-third of the global population (Sanchez and Besteiro, 2021). Due to emerging resistance and potential side effects of some existing treatments, new and improved medicines are always needed to cure apicomplexan-caused diseases.

The apicoplast is a validated drug target, not only because it hosts crucial housekeeping metabolic pathways, but also because the proteins involved in these pathways are essentially of prokaryotic origin, which may facilitate the design of inhibitors with high specificity. The main metabolic pathways found in the organelle are heme synthesis (jointly with the mitochondrion), the type II fatty acid synthesis (FASII) pathway, the synthesis of isoprenoids (which are important lipid precursors), and the synthesis of iron-sulfur (Fe-S) clusters, which is central because these cofactors are essential for enzymes involved in the other apicoplast-hosted pathways (Seeber and Soldati-Favre, 2010). Not all apicoplast-based pathways may have the same importance in all species or developmental stages, but they contribute to general parasite fitness. In the case of the tachyzoite forms of *T. gondii* (highly multiplicative forms of the parasite, responsible for the acute phase of the disease), the organelle is clearly essential for parasite growth. Contrarily to *P. falciparum* blood stages that mainly utilizes host derived heme, in *T. gondii* tachyzoites are unable to efficiently salvage heme for which de novo synthesis is thus important (Bergmann *et al*., 2020). FAS II is also contributing significantly to parasite fitness (Mazumdar *et al*., 2006), although scavenging of FA precursors from the host cells can at least partially compensate for lack of *de novo* synthesis (Amiar *et al*., 2020; Liang *et al*., 2020). Isoprenoid synthesis, which is in fact the only essential apicoplast-hosted pathway in *Plasmodium* blood stages (Yeh and DeRisi, 2011) and one of the most conserved function of the apicoplast in Apicomplexa, is also clearly essential for the growth of *T. gondii* tachyzoites (Nair *et al*., 2011). Finally, synthesis of Fe-S clusters in the apicoplast, which provides important cofactors for proteins that are in turn involved in fitness-conferring pathways of the organelle like FASII or isoprenoid synthesis, is essential for the viability of the parasites (Pamukcu *et al*., 2021; Renaud *et al*., 2022).

In order to function properly, these pathways must be supplied with key precursors originating from other organelles or from the cytoplasm and probably imported through specific transporters (Kloehn *et al*., 2021). Among them carbon precursors for the synthesis of FAs and isoprenoids are imported in the form of pyruvate, as well as glycolytic intermediates glyceraldehyde 3-phosphate (GAP) and dihydroxyacetone phosphate (DHAP). These glycolytic intermediates are imported into the organelle from the cytosol (Brooks *et al*., 2010; Chen *et al*., 2024). Apicoplast-specific isoforms of glycolytic enzymes such as triosephosphate isomerase, GAP dehydrogenase and pyruvate kinase and phosphoglycerate kinase are both involved in the production of ATP and reducing power in the organelle, but also feed the FASII (through the synthesis of acetyl-CoA from pyruvate) as well as isoprenoid synthesis (through the synthesis of 1-deoxy-D-xylulose 5-phosphate from pyruvate and DHAP) (Niu *et al*., 2022). GAP and DHAP, three-carbon compounds that can be interconverted, are thus key to apicoplast metabolism, however they also be source of toxicity as they can convert spontaneously into methylglyoxal (MG), a highly reactive dicarbonyl compound that is involved in the formation of advanced glycation end products (AGEs) (Allaman *et al*., 2015). The glycation of proteins, nucleic acids and lipids is detrimental to their function and, more generally, to the cell. Glycolysis is thus usually accompanied by a cytosolic glutathione (GSH)-dependent methylglyoxal (MG) detoxification system made of two sequential reaction mediated by glyoxalase 1 (Glo1) and glyoxalase 2 (Glo2) enzymes, that will convert MG to to D-lactate (Morgenstern *et al*., 2020). More precisely, MG spontaneously reacts with GSH to form a hemithioacetal adduct which is then metabolized by Glo1, a metalloenzyme requiring divalent metal ions (either Ni^2+^ or Zn^2+^) for activation, to form S-lactoylglutathione. Subsequently, Glo2 converts S-lactoylglutathione to D-lactate, recycling GSH in the process. A more direct route for MG detoxification also exists in some organisms, which involves glyoxalase 3 (Glo3) to catalyze the conversion of MG to D-lactate in one step (without involving GSH or metal ions) (Misra *et al*., 1995).

In addition to tricarboxylic acid cycle, apicomplexan parasites like *P. falciparum* and *T. gondii* use glycolysis for energy production (Jacot *et al*., 2016). Accordingly, these parasites contain putative homologues of the Glo1 and Glo2 enzymes (Deponte, 2014). However, they have mostly been studied in *P. falciparum*, where investigations identified a cytosolic Glo1/Glo2 system (Akoachere *et al*., 2005; Urscher *et al*., 2011), which was later found to be not essential for the viability of blood stages of the parasites (Wezena *et al*., 2017). Similarly, the *T. gondii* homologues of Glo1 and Glo2 were predicted to be dispensable by a genome-wide CRISPR-based phenotypic screen (Sidik *et al*., 2016). Interestingly, an apicoplast-localized Glo2 isoform in *P. falciparum*, as well as a Glo1-like protein (PfGilp) also targeted to the organelle were identified (Akoachere *et al*., 2005). This suggested the presence of an apicoplast-specific MG detoxification system, in line with the presence of GAP and DHAP in the organelle, however PfGilp activity could not be obtained *in vitro*, possibly because some active site residues are missing (Akoachere *et al*., 2005). A pathway that would be specifically dedicated to MG detoxification in the organelle in Apicomplexa is thus possibly present, but it has not yet been shown to be functional.

In *T. gondii*, where MG detoxification is much less characterized than in *P. falciparum*, we have identified an unusual Glo1-motif containing protein predicted to localize to the apicoplast. We present here the functional characterization of this protein and show that it is essential for maintaining the homeostasis of the organelle.

## Results

### TgGloL is an apicoplast-localized Glo1 domain-containing protein

Using homology searches, we looked for potential Glo1 or Glo2 homologues in *T. gondii* homologues. A Glo1 isoform was already confirmed to localize to the cytosol (Goo *et al*., 2015), although it was not really characterized further, and is thus likely a component of the cytosolic MG-detoxification system (Fig. 1A). As for the *P. falciparum* homologue, the protein contains two Glo1 domains. Glo1 belongs to the vicinal oxygen chelate (VOC) metalloenzyme superfamily whose members contain a paired βαβββ motif involved in metal coordination (He and Moran, 2011). While Glo1 enzymes typically function as homodimers in some organisms, including Apicomplexa, they are monomeric containing two Glo1 domains encoded by a single polypeptide (Iozef *et al*., 2003). Contrarily to *P. falciparum* that expresses both a cytosolic an apicoplast-localized isoform of Glo2, a single Glo2 homologue was found for *T. gondii*, with no predicted transit peptide for targeting to the organelle (Fig. 1A). Interestingly, besides TgGlo1 we also found an atypical protein containing Glo1-like motifs potentially encoded by the *T. gondii* genome (Fig. 1A, entry TGGT1_240430 in the www.Toxodb.org database). This large protein did not have any particular motif beyond the Glo1 domains, but CRISPR-based data suggested it might be essential (Sidik *et al*., 2016) and data from global mapping of *T. gondii* proteins subcellular location by spatial proteomics (Barylyuk *et al*., 2020) suggested an apicoplast localization. However, this protein (that we called TgGloL for “glyoxalase-like”) did not seem to bear significant homology with the apicoplast-localized *P. falciparum* GILP. Accordingly, the phylogenetic analysis of members of the Glo1/Glo2 families we performed positioned TgGloL close to Glo1 cluster, but clearly as an outlier and also clearly distinct from GILP (Fig. 1B).

**Figure 1.**
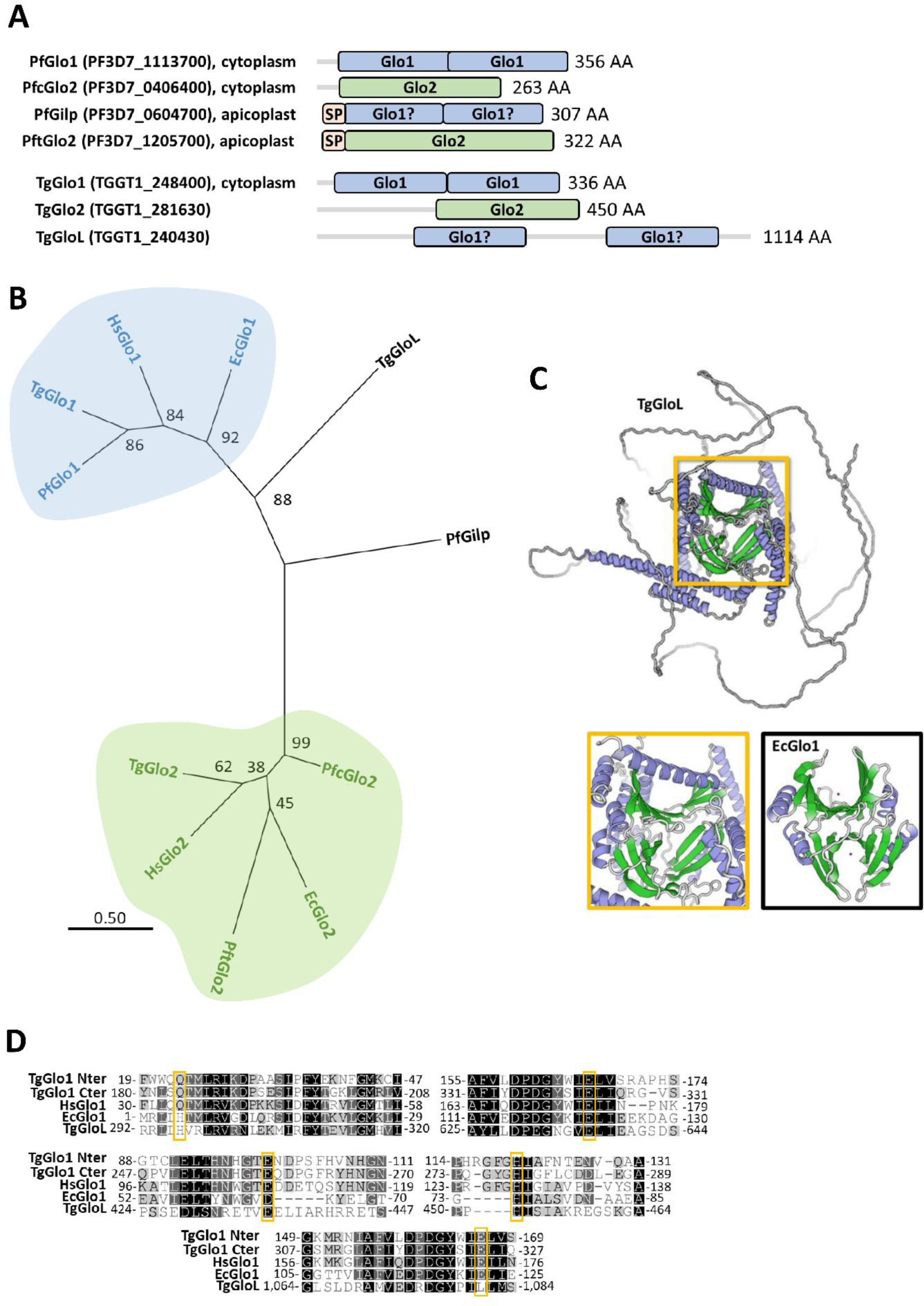
TgGloL is an atypical glyoxalase domain-containing protein. **A)** Organization of glyoxalase 1 (Glo1) and 2 (Glo2) domain-containing proteins encoded by *Plasmodium falciparum* (Pf) and *Toxoplasma gondii* (Tg). **B)** Phylogenetic analysis of glyoxalase domain-containing proteins of Pf and Tg, as well as *Homo sapiens* (Hs) and *Escherichia coli* (Ec) Glo1 and Glo2 sequences. The percentage of replicate trees in which the associated taxa clustered together in the bootstrap test 500s replicates are shown next to the branches (bootstrap value) and cale bar with amino acid substitutions per site is also displayed. **C)** Homology-derived structural model of TgGloL (top) and comparison (bottom) of the structured core of TgGloL with the βαβββ motif of EcGlo1 (protein data bank entry: 1F9Z). **D)** CLUSTAL alignments of selected regions of TgGloL as well as Glo1 from *T. gondii* (both N-terminal and C-terminal domains), *H. sapiens* and *E. coli*, showing various degrees of conservation of residues potentially involved in enzyme activity (squared).

Structural modelling with the deep-learning AlphaFold2 algorithm (Jumper *et al*., 2021) revealed that besides long unstructured loops, TgGloL has a well-structured core whose organization resembles the double association of βαβββ Glo1 motifs typically found in the experimentally-determined 3D structure of the *Escherichia coli* Glo1 homodimer for example (Fig. 1C). Analysis of the TgGloL primary amino acid sequence by aligning and comparing with sequences from different Glo1 homologues, we could see that while several, but not all, residues potentially involved in the coordination of the metal ion (He and Moran, 2011) were conserved (Fig. 1D). As for GILP, this raises a doubt about the ability of this protein to act as an active Glo1 enzyme.

To localize TgGloL, we epitope-tagged the native protein. This was achieved in the TATi-ΔKu80 cell line, which not only favors homologous recombination, but would also allow generating an anhydrotetracycline (ATc)-regulatable conditional mutant. A sequence coding for a C-terminal triple hemagglutinin (HA) epitope tag was inserted at the endogenous *TgGLoL* locus by single homologous recombination (Fig. 2A). Transfected TATi-ΔKu80 parasites were subjected to chloramphenicol selection and clones were isolated and checked for correct integration by PCR (Fig. 2B). The resulting cell line, named TgGloL-HA, was then checked by immunoblot with an anti-HA antibody, which labelled two main products (Fig. 2C), at around the predicted size for TgGloL (123 kDa). This would be consistent with precursor and mature forms of TgGloL, that would be the result from the cleavage of the transit peptide upon import into the apicoplast. Immunofluorescence assay (IFA) with anti-HA antibody and co-staining with an apicoplast marker confirmed that TgGloL localizes to the organelle (Fig. 2D).

**Figure 2.**
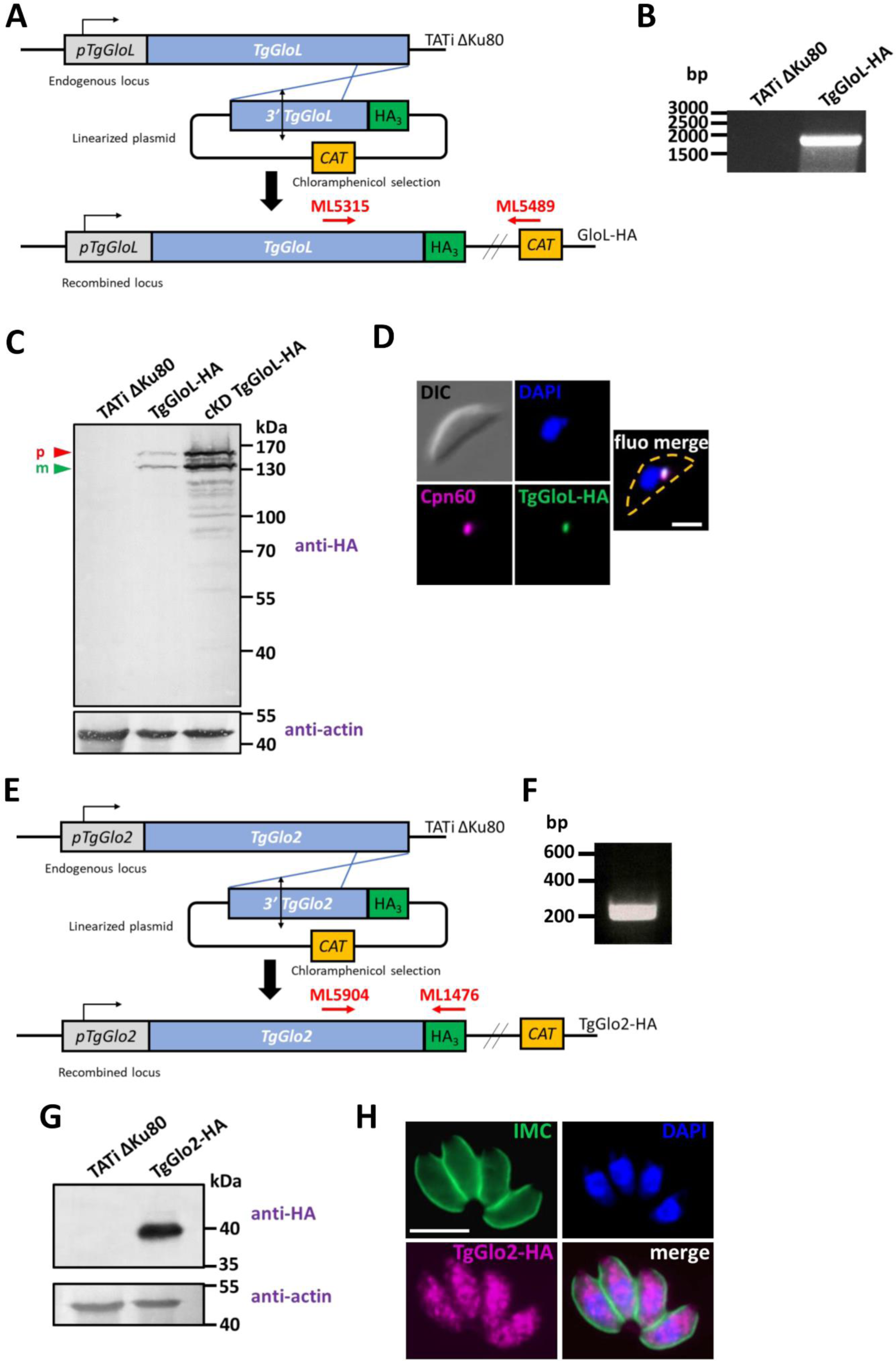
TgGloL localizes to the apicoplast while TgGlo2 is cytosolic. **A)** Schematic representation of the strategy for expressing HA-tagged version of TgGloL by homologous recombination at the native locus of the corresponding gene of interest. Chloramphenicol was used to select transgenic parasites based on their expression of the Chloramphenicol acetyltransferase (CAT). **B)** Diagnostic PCR for verifying correct integration of the construct. The amplified fragment corresponds to the red arrows in **A)**, and specific primers used were ML5315/ML5489. **C)** Immunoblot analysis with anti-HA antibody shows precursor (p) and mature (m) forms of C-terminally HA-tagged TgGloL in TgGloL-HA and cKD TgGloL-HA cell lines. **D)** HA-tagged TgGloL (green) localizes to the apicoplast (co-stained with marker TgCpn60, magenta). Scale bar represents 5 μm. DNA was labelled with DAPI. DIC: differential interference contrast. Parasite shape is outlined on merged image. **E)** Schematic representation of the strategy for expressing HA-tagged version of TgGlo2 by homologous recombination at the native locus of the corresponding gene of interest. Chloramphenicol was used to select transgenic parasites based on their expression of the CAT. **F)** Diagnostic PCR for verifying correct integration of the construct. The amplified fragment corresponds to the red arrows in **E)**, and specific primers used were ML5904/ML1476. **C)** Immunoblot analysis with anti-HA antibody shows a single product at around 40 kDa. **D)** HA-tagged TgGlo2 (magenta) localizes to the cytosol, in parasites delineated by a TgIMC3 labelling (green). Scale bar represents 5 μm. DNA was labelled with DAPI.

As the only TgGlo2 isoform encoded by the genome of *T. gondii* (entry TGGT1_281630 in the www.Toxodb.org database) had never been properly localized, we also epitope-tagged this protein using a similar strategy (Fig. 2E). A selected clone was check by PCR (Fig. 2F) and in sharp contrast to TgGloL, immunoblot analysis revealed a single product for this protein (Fig. 2E) and IFA showed a cytoplasmic distribution of the protein. This suggests that TgGlo2 is likely part of the cytosolic MG detoxification pathway and is not associated with the apicoplast.

### TgGloL is important for parasite growth

Next, we generated a conditional TgGLoL mutant cell line in the TgGloL-HA–expressing TATi ΔKu80 background. Replacement of the endogenous promoter by an inducible-Tet07SAG4 promoter was achieved through a single homologous recombination at the locus of interest, yielding the conditional knockdown (cKD) TgGloL-HA cell line (Fig. 3A). Integration was verified by PCR (Fig. 3B) and immunoblot analysis showed that the promoter change led to an increase in protein expression over the native TgGloL promoter (Fig. 2C). Immunoblot analysis nevertheless confirmed that incubation of the cell line with ATc efficiently down-regulated TgGloL expression levels, although it necessitated several days of incubation (Fig. 3C). IFA also confirmed the down-regulation of the protein upon ATc treatment, as it became almost undetectable four days post-treatment (Fig. 3D). This effect was specific to TgGloL, because in these conditions the Cpn60 apicoplast marker was not affected (Fig. 3D).

**Figure 3.**
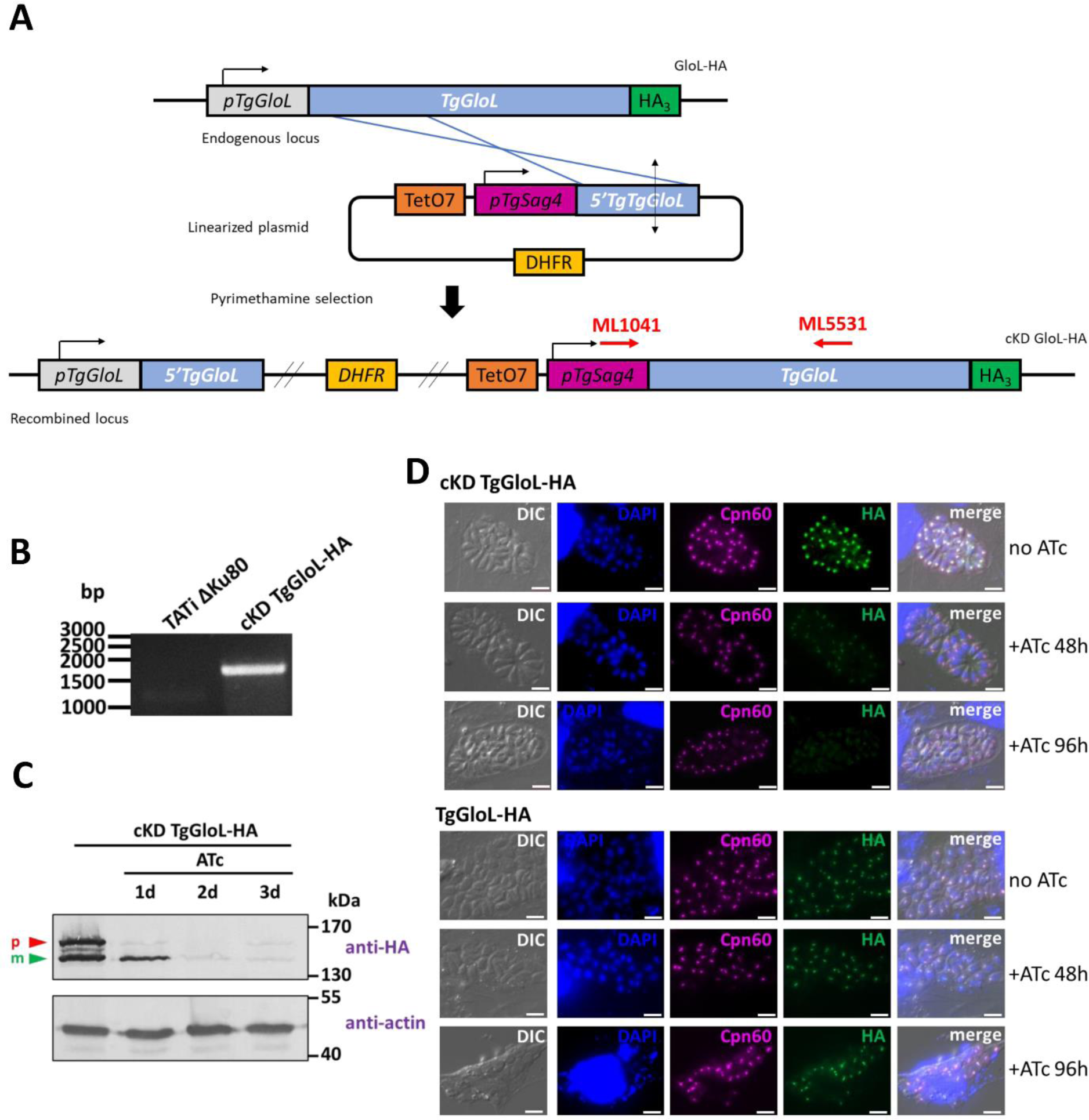
Generating a conditional knock-down mutant for TgGloL. **A)** Schematic representation of the strategy for generating TgGloL conditional knock-down cell line by homologous recombination at the native locus. Pyrimethamine was used to select transgenic parasites based on the expression of dihydrofolate reductase (DHFR). **B)** Diagnostic PCR for verifying correct integration of the construct. The amplified fragments confirming integration, corresponding to the red arrows displayed in **A)**, and specific primers used were ML1041/ML5531. **C)** Immunoblot analysis with anti-HA antibody shows precursor (p) and mature (m) forms of C-terminally HA-tagged TgGloL and efficient down-regulation of the protein after 48 hours of incubation with anhydrotetracycline (ATc). Anti-actin antibody was used as a loading control. **D)** Immunofluorescence assay confirms down-regulation of HA-tagged TgGloL (green) from 48 hours and up to 96 hours of ATc incubation, with little impact on the TgCpn60-labelled apicoplast. In these experimental conditions ATc had no impact on TgGloL or Cpn60 signals in the parental TgGloL-HA cell line. Scale bar represents 5 μm. DNA was labelled with DAPI. DIC: differential interference contrast.

We then assessed the impact of TgGloL depletion on *in vitro* parasite fitness by performing a plaque assay, which determines the ability of cKD TgGloL-HA parasites to generate lysis plaques on a host cell monolayer in the absence or continuous presence of ATc for 7 days. TgGloL depletion largely prevented plaque formation, indicating that the protein is important for parasite fitness (Fig. 4A, B). In order to assess if the impact on the parasites was sublethal, we performed an experiment where ATc was washed out (or not) after 7 days of incubation and monitoring plaque formation after another 7 days of incubation. We could observe some partial recovery after ATc washout, indicating that not all parasites died after 7 days of TgGloL depletion (Fig. 4C, D).

**Figure 4.**
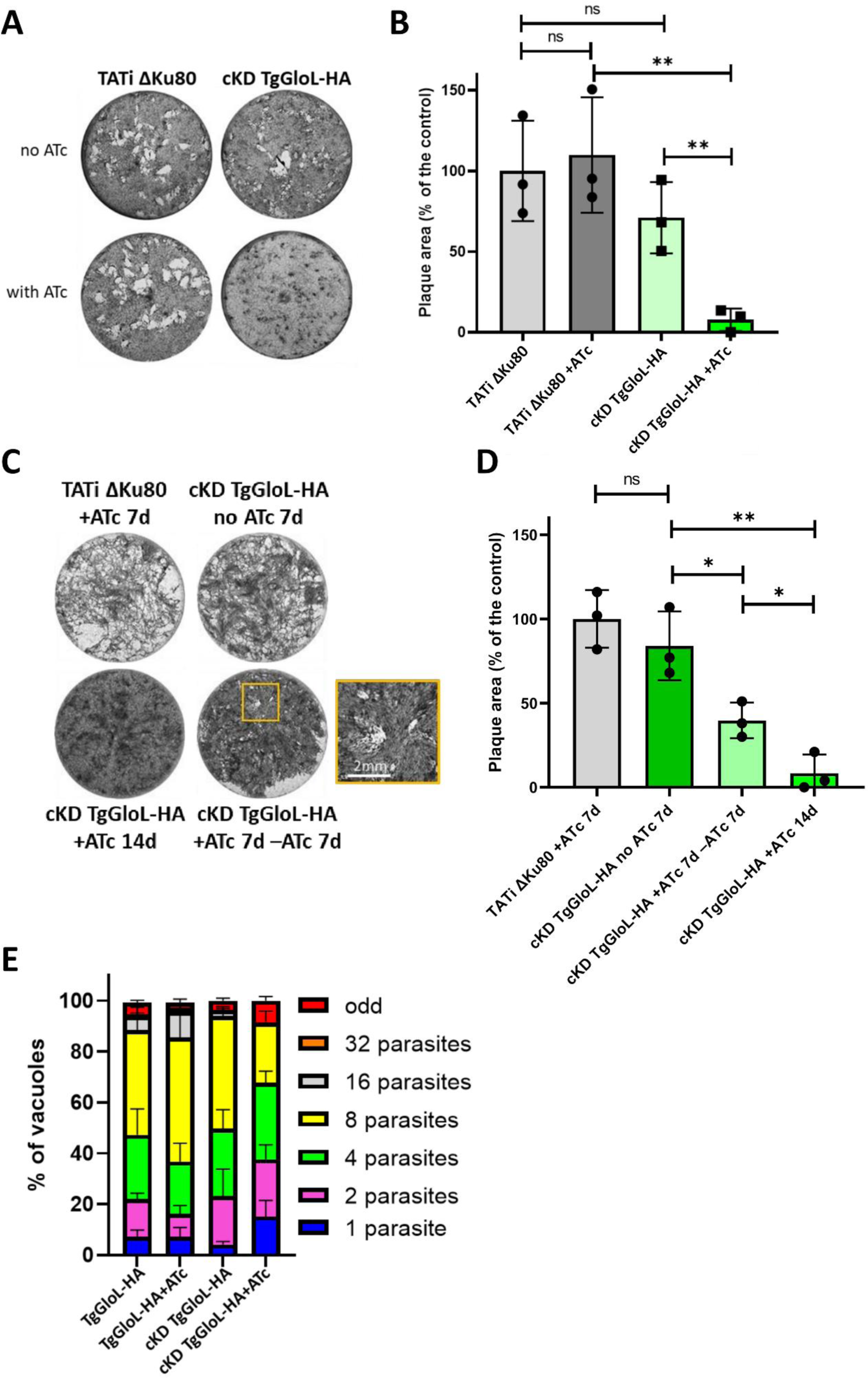
TgGloL is important for parasite fitness. **A)** Plaque assays were carried out by infecting HFF monolayers with the cKD TgGloL-HA cell line or TgGloL-HA control. They were grown for 7 days ± ATc. **B)** Measurements of lysis plaque areas highlight a significant defect in the lytic cycle when TgGloL is depleted. Values are means of n = 3 experiments ± SD. Mean value of TgGloL-HA control grown in the presence of ATc was set to 100% as a reference. **, p≤0.01; ns, not significant; two-tailed Student’s t-test p-values. **C)** Plaque assays for the TgGloL mutant was performed as described in A), but ATc was washed out after 7 days (+ATc 7d -ATc 7d) or not (+ATc 14d), and parasites were left to grow for an extra 7 days. Small plaques were observed upon ATc removal (inset), suggesting some parasites were still viable after 7 days of treatment. Shown are images from one representative out of three independent experiments. **D)** Measurements of lysis plaque areas from the washout experiment described in **C)**. Values are means of n = 3 experiments ± SD. Mean value of TgGloL-HA control grown in the presence of ATc for 7 days was set to 100% as a reference. *, p≤0.05; **, p≤0.01; ns, not significant; two-tailed Student’s t-test p-values. **E)** The cKD TgGloL-HA parasites, as well as the TgGloL-HA control, were grown in HFF in the presence or absence of ATc for 48 hours, and subsequently allowed to invade and grow in new HFF cells for an extra 24 hours in the presence of ATc. Parasites per vacuole were then counted. Values are means ± SD from n = 3 independent experiments for which 200 vacuoles were counted for each condition.

Lysis plaques are the result of successive cycles of invasion, replication and egress by the parasites on the host cell monolayer. To assess if the replication step was particularly affected, we thus performed a replication assay. The cKD TgGloL cell line was preincubated in the absence or presence of ATc for 48h and released mechanically, before infecting new host cells and were then grown for an additional 24h in ATc prior to parasite counting. We noted that the incubation with ATc led to an accumulation of vacuoles with fewer TgGloL mutant parasites, indicating that the protein is important for parasite replication (Fig. 4E).

### TgGloL is important for maintaining apicoplast homeostasis

To get more insights on the impact of TgGLoL depletion at the subcellular level, we then performed a morphological analysis by electron microscopy. Parasites were incubated for 5 days in the presence of ATc to efficiently deplete TgGloL, and then left to invade new host cells and incubated for 2 extra days in the presence of ATc prior to sample processing and image acquisition. We observed some important organellar segregation defects during cell division (Fig. 5A), as well as severe membrane-related problems, with discontinuous vesicular material trapped between divided parasites, instead of a continuous plasma membrane (Fig. 5A, B). Moreover, while apicoplasts were rarely detected upon TgGloL detection, the membranes of those that could be observed appeared loosened, and the lumen of the organelle appeared less dense and even electron lucent, suggesting an alteration of their content (Fig. 5B). These observations were in sharp contrast with the aspect of the apicoplast or the plasma membrane in the TgGloL-expressing control (Fig. 5C). Interestingly, these phenotypic manifestations were very similar to those observed in mutants of the apicoplast-localized Fe-S assembly pathway in which important metabolic functions of the organelle are affected (Renaud *et al*., 2022).

**Figure 5.**
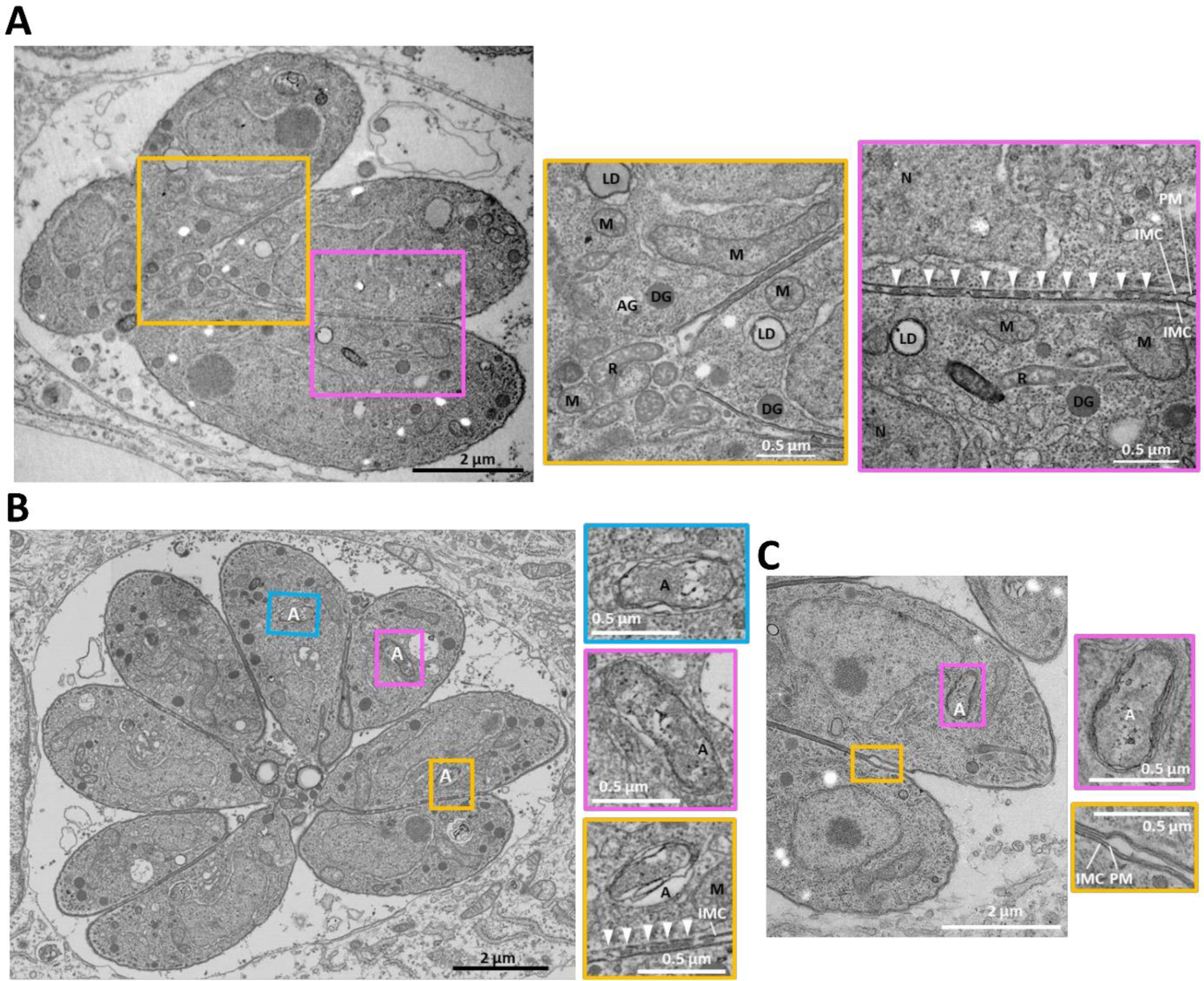
Electron microscopy analysis of cKD TgGloL-HA parasites grown for up to 7 days in the presence of ATc shows several ultrastructural defects. **A)** TgGloL depletion leads to default in plasma membrane separation during parasite division, with trapped material between unseparated parasites (arrowheads), as displayed on insets representing magnifications of selected parts of the respective left image. **B)** TgGloL depletion also alters the aspect of the apicoplast (loosened membranes and less dense content) in parasites in which the organelle is still visible (insets show magnification of selected organelles). **C)** Aspect of the plasma membrane (yellow inset) and apicoplast (magenta inset) in control parasites. AG: amylopectin granule, DG: dense granule, IMC: inner membrane complex, LD: lipid droplet, M: mitochondrion, PM: plasma membrane, R: rhoptry.

To further assess the specific impact of TgGloL depletion on the apicoplast, we performed IFA on cKD TgGloL parasites grown for extended periods of time in the presence of ATc. We did not observe any drastic morphological alteration of the mitochondrion (typically present as a single lasso-shaped network in intracellular parasites) or micronemes (secretory organelles located at the apex of the parasites) even after 7 days of ATc treatment (Fig. 6A, C). In sharp contrast the Cpn60 apicoplast marker, which appeared essentially unaltered after 4 days of ATc treatment (Fig. 3D), appeared less evenly distributed between parasites after 5 days of ATc treatment, and largely absent from the parasites after 7 days (Fig. 6A). To confirm this was not due to a specific impact on this protein, but a more general organellar defect, we also labelled the E2 subunit of succinate dehydrogenase (SDH), another apicoplast marker and observed a similar trend (Fig. 6C). In both cases, quantification confirmed that up to 80% of the parasites had likely lost the organelle after 7 days of ATc treatment, which was not observed when the control cell line was treated with ATc for the same duration (Fig. 6B, D).

**Figure 6.**
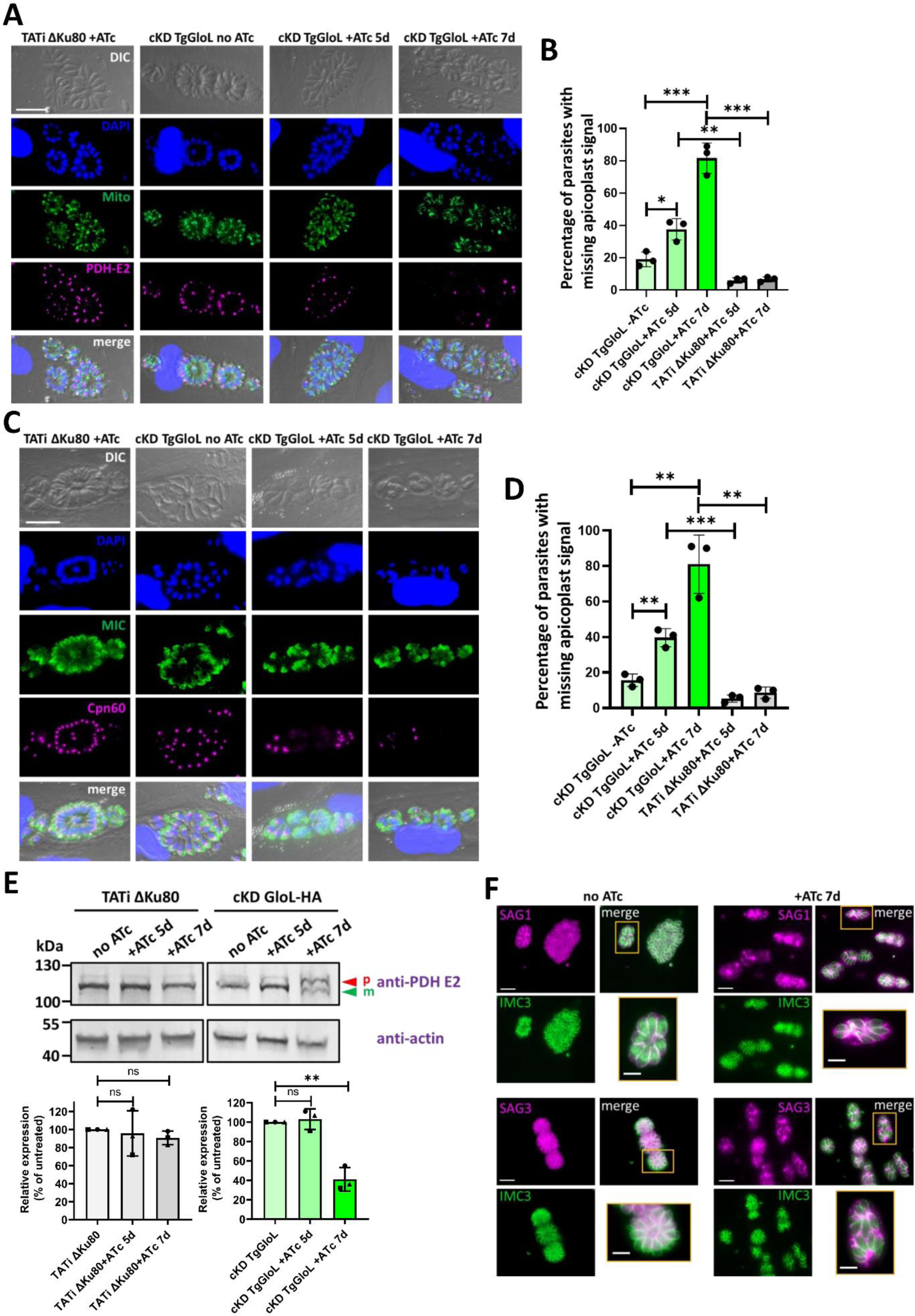
Long term TgGloL depletion leads to a loss of the apicoplast. **A)** cKD TgGloL parasites were treated for 5 or 7 days with ATc and stained for the mitochondrion or the apicoplast (with anti-E2 subunit of the pyruvate dehydrogenase -PDH-). TATi ΔKu80 parasites treated for 7 days with ATc were used as a control to assess a potential impact of ATc on the organelles. Scale bar represents 10 μm. DNA was labelled with DAPI. DIC: differential interference contrast. **B)** Quantification of PDH-E2 signal loss in cKD TgGloL parasites were treated for 5 or 7 days. Data are mean ± SD from *n*=3 independent experiments. *, p≤0.05; **, p≤0.01; ***, p≤0.001; two-tailed Student’s t-test p-values. **C)** cKD TgGloL parasites were treated for 5 or 7 days with ATc and stained with a microneme marker or for the apicoplast (with anti-Cpn60). TATi ΔKu80 parasites treated for 7 days with ATc were used as a control to assess a potential impact of ATc on the organelles. Scale bar represents 10 μm. DNA was labelled with DAPI. DIC: differential interference contrast. **D)** Quantification of Cpn60 signal loss in cKD TgGloL parasites were treated for 5 or 7 days. Data are mean ± SD from *n*=3 independent experiments. **, p≤0.01; ***, p≤0.01; two-tailed Student’s t-test p-values. **E)** Immunoblot analysis (top) and quantification (bottom) of the mature form (m) of PDH-E2 that decreases over time in the cKD TgGloL cell line upon treatment with ATc, while the precursor form accumulates (p). Quantifications were performed from *n*=3 independent experiments and are represented as mean ± SD. **, p≤0.01; ns, not significant; two-tailed Student’s t-test p-values. **F)** cKD TgGloL parasites mutants were grown for up to 7 days in the presence or absence of ATc and then co-stained for inner membrane complex marker IMC3 (green) together with GPI-anchored protein SAG3 (magenta). As shown on insets representing selected parts of the images, the depletion of TgGloL leads to the accumulation of SAG3 in the vacuolar space. Scale bar represents 5 μm. DNA was labelled with DAPI. DIC: differential interference contrast.

Immunoblot analysis of the lumen apicoplast protein PDH E2 upon TgGloL depletion confirmed a decrease in the abundance of its mature form over the precursor form (Fig. 6E). Again, this would be consistent with a loss of the organelle, as accumulation of unprocessed protein is typically used as a marker for the loss of protein import that accompanies apicoplast loss (Yeh and DeRisi, 2011). As one of the main function of the apicoplast is to generate isoprenoid precursors which are central to a number of important posttranslational protein modifications like glycosylphosphatidylinositol (GPI) anchoring (Imlay and Odom, 2014), we assessed the localization of GPI-anchored surface antigen (SAG) proteins SAG1 and SAG3 in TgGloL-depleted parasites. We observed obvious signs of mislocalization for these two proteins: instead of being homogenously distributed at the periphery of the parasites, they were often seen accumulated in patches or even found within the parasitophorous vacuole space (Fig. 6F). Again, this is reminiscent of the phenotype observed for mutants of the apicoplast-localized Fe-S assembly pathway that are defective for apicoplast function (Renaud *et al*., 2022).

### No evidence that MG plays a role in the demise of TgGloL mutant parasites

We wanted to assess if TgGloL could act as a Glo1 enzyme in spite of its noticeable differences in its putative active site (Fig. 1D). Unfortunately, we failed to express a recombinant version of the protein in *E. coli*, in spite of multiple attempts to modulate expression conditions. Alternatively, we sought to detect a potential accumulation of MG or AGEs upon TgGloL depletion. MG is highly reactive and AGEs are often in low abundance and are very heterogenous, which renders their detection by proteomics or metabolomics approaches particularly challenging and prone to inconsistencies (Donnellan *et al*., 2022). However, specific anti-AGE antibodies have been used with some success in the past in *P. falciparum* Glo mutants (Wezena *et al*., 2017). We thus used two commercial monoclonal antibodies raised against different MG-modified carrier proteins to detect AGEs in the presence or absence of TgGloL. By immunoblot, the two antibodies revealed similar complex profiles with staining patterns showing two main products at around 35 and 55 kDa in addition to several less-well detected products (Fig. 7A). Importantly, there was no particular change in the profiles or in abundance of some products upon depletion of TgGloL (Fig. 7A). We next used these antibodies to check for a potential accumulation of AGE in the apicoplast upon TgGloL depletion. We performed IFA on cKD TgGloL-HA parasites incubated for 5 days with ATc, as not all parasites have lost the organelle at this point (Fig. 6A-D). Both anti-AGE antibodies stained puncta in the host cells and the parasites, but co-staining with an apicoplast marker did not reveal any specific co-localization between the AGE signal and the apicoplast (Fig. 7B). This suggests that AGEs are not accumulating in the organelle when TgGloL is depleted.

**Figure 7.**
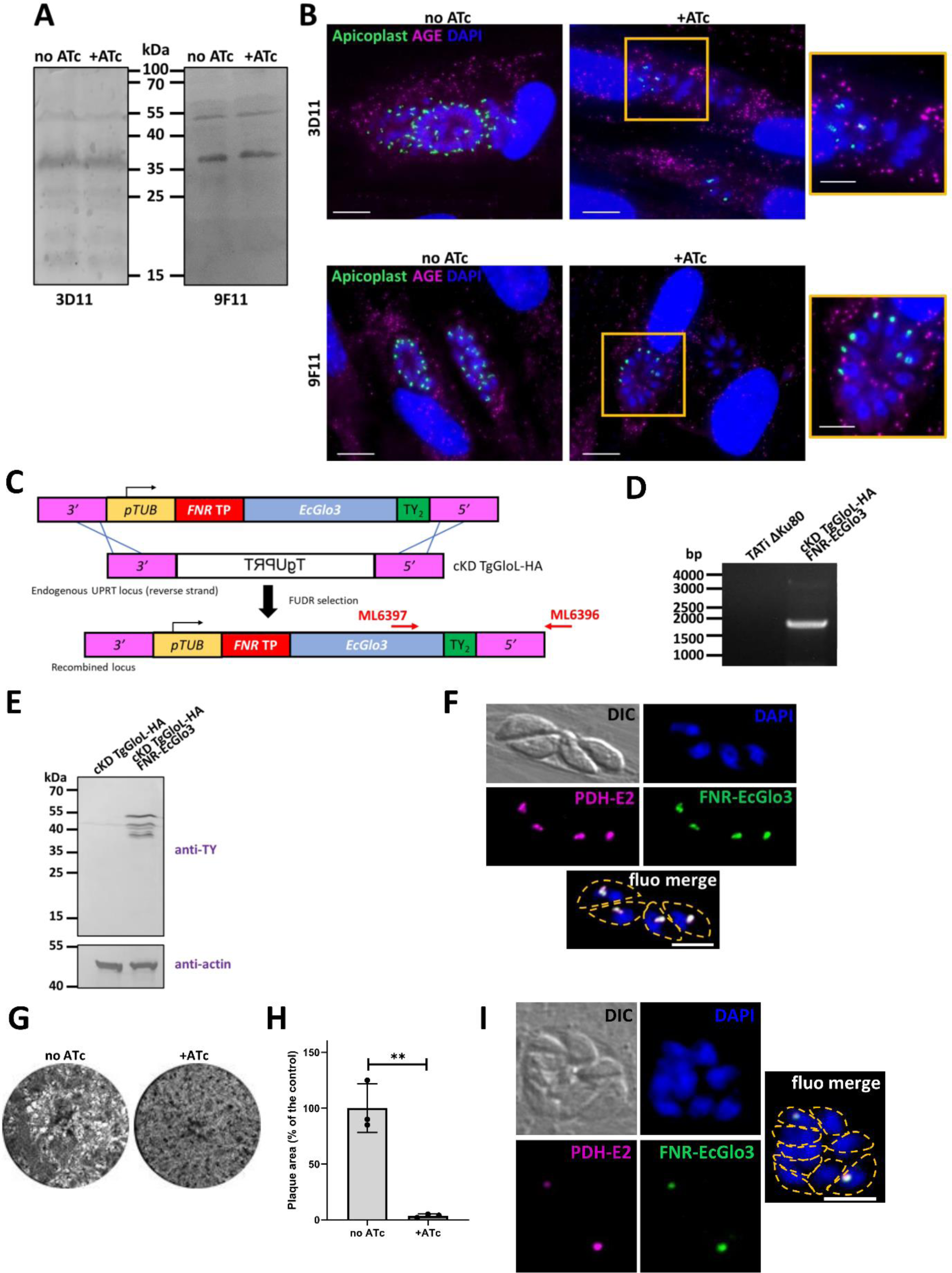
No evidence for methylglyoxal accumulation in the apicoplast upon TgGloL depletion. A) Immunoblot analysis of cKD TgGloL parasites treated for 5 days with ATc or not with two different monoclonal antibodies (3D11 and 9F11) against glycated products does not show particular quantitative or qualitative differences induced by TgGloL depletion. **B)** The same antibodies (magenta) failed to detect a signal in the apicoplast (stained with anti-PDH-E2, green) by immunofluorescence upon TgGloL depletion. Scale bar represents 10 µm or 5 μm (insets). DNA was labelled with DAPI. **C)** Schematic representation of the strategy for expressing apicoplast-targeted *E. coli* glyoxalase 3 (EcGlo3) by homologous recombination at the uracil phosphoribosyl transferase (UPRT) locus, allowing for selection of transgenic parasites with 5-fluorodeoxyuridine (FUDR). **D)** Diagnostic PCR for verifying correct integration of the construct. The amplified fragments confirming integration, corresponding to the red arrows displayed in **C)**, and specific primers used were ML6396/ML6397. **E)** Immunoblot analysis shows precursor (p) and mature (m) forms of C-terminally TY-tagged FNR-EcGlo3 expressed in the cKD TgGloL-HA background. **F)** TY-tagged FNR-EcGlo3 (green) localizes to the apicoplast (co-stained with marker PDH-E2, magenta). Scale bar represents 5 μm. DNA was labelled with DAPI. DIC: differential interference contrast. Parasite shape is outlined on merged image. **G)** Representative plaque assay (out of 3 independent biological replicates) performed in the cKD TgGloL-HA cell line complemented with apicoplast localized EcGlo3 showing absence of compensation for the fitness defect. **H)** Measurements of lysis plaque areas in cKD TgGloL-HA FNR-EcGlo3 parasites grown in the presence of ATc or not for 7 days. Values are means of n = 3 experiments ± SD. Mean value of the cell line grown in absence of ATc was set to 100% as a reference. **, p≤0.01; two-tailed Student’s t-test p-value. **I)** immunostaining of the apicoplast showing organelle loss in the cKD TgGloL-HA FNR-EcGlo3 cell line kept for 7 days in the presence of ATc (representative image was chosen out of 2 independent biological replicates). Scale bar represents 5 μm. DNA was labelled with DAPI. DIC: differential interference contrast. Parasite shape is outlined on merged image.

It is possible that MG or AGE are potentially present in harmful concentration in the apicoplast upon TgGloL depletion, but that immunochemistry-based approaches are not sensitive or specific enough to detect an accumulation. We thus reasoned that if this is the case, expressing a MG-detoxifying enzyme in the organelle may help restoring fitness in TgGloL-depleted parasites. *E. coli* Glo3 can potentially catalyze the conversion of MG to D-lactate in one step (Misra *et al*., 1995), so we constructed a chimeric version of EcGlo3 fused to a N-terminal apicoplast-targeting sequence (the transit peptide from the ferredoxin-NADP^+^ reductase -FNR-(Harb *et al*., 2004)), to be stably integrated at the non-essential *uracil phosphoribosyl transferase* locus in the TgGloL conditional mutant (Fig. 7C, D). This protein was tagged with two TY epitopes (Bastin *et al*., 1996) and detection by immunoblot revealed multiple specific products at around 50 kDa (Fig. 7E). This is likely to reflect the proteolytic maturation of the protein in the lumen of the apicoplast and is in accordance with the expected size of the full-length fusion protein, which is 46 kDa (36 kDa after the removal of the transit peptide sequence). IFA showed that TY-tagged FNR-EcGlo3 is co-localizing with the apicoplast lumen marker TgPDH-E2 (Fig. 7F). We next performed plaque assays in the presence of ATc to see if the expression of apicoplast-localized EcGlo3 could compensate for the depletion of TgGloL, and at least partially restore parasite fitness (Fig. 7G, H). We observed no obvious restoration of fitness upon depletion of TgGloL in spite of the expression of EcGlo3 in the apicoplast. Moreover, apicoplast labelling in parasites kept in the presence of ATc for up to 7 days revealed a marked impact on the organelle in this cell line, hinting that the expression of EcGlo3 was not sufficient to restore the long-term detrimental effects of TgGloL depletion on the organelle (Fig. 7I). Of course, we have no evidence that the apicoplast-expressed EcGlo3 is fully functional or expressed at sufficient levels to efficiently detoxify MG locally, but overall our data does not point to the accumulation of MG as being the cause for the demise of TgGloL-depleted parasites.

## Discussion

Even in normal homeostatic growth conditions cells may generate toxic endogenous molecules through their metabolic activity. These may be molecules that can act as competitive analogs against other metabolites, or molecules with highly reactive groups that allow them to covalently modify cellular components and thus impair their function. Although some enzymes have evolved to convert these toxic metabolites into less reactive molecules that may be easily cleared or recycled, there is a fine balance between toxic metabolite production and detoxification that needs to be maintained to prevent cellular damage (Lee *et al*., 2020).

MG is a reactive aldehyde, mostly generated through glycolysis (Allaman *et al*., 2015), that spontaneously reacts with proteins, lipids and nucleic acids to form toxic AGEs. Specific systems for sensing and detoxifying MG are present in all domains of life, including eukaryotes, but also ancient archae (Oren and Gurevich, 1995) and bacteria (Lee and Park, 2017), as even prokaryotic species not relying on glycolysis but that use alternative pathways instead for central metabolism are known to synthesize some MG. During evolution, the MG-detoxifying systems have undergone significant evolutionary changes and diversification, likely to confer some flexibility and adaptation in their functions: besides the two-enzyme GSH-dependent systems with Glo1/Glo2, there is also for instance the GSH-independent single Glo3 enzyme (Misra *et al*., 1995; Kumar *et al*., 2021).

The acquisition of an apicoplast in Apicomplexa species has brought important metabolic functions to these parasites, one of the most (if not the most) important is the synthesis of isoprenoid precursors, but this necessitates the import of MG-generating molecules DHAP and GAP into the organelle (Fig. 8). How the toxic potential of these glycolytic intermediates is alleviated is currently unknown. Although the Glo-based systems are the main route for MG detoxification in living cells, the possibility that MG-derivatives could be exported by efflux pumps has been recently suggested (Coukos and Moellering, 2021). The export of MG from the organelle to be detoxified by the Glo1/Glo2 system expressed in the cytosol of apicomplexan parasites could thus be an alternative pathway to local detoxification (Fig. 8), but the complement of apicoplast-localized transporters is currently poorly characterized (Kloehn *et al*., 2021) and no such candidate has yet been identified. Moreover, in plastids like the chloroplast, in which MG is produced as a byproduct of the Calvin cycle, Glo-based systems seem to be the main detoxification pathways (Shimakawa *et al*., 2018; Kumar *et al*., 2021).

**Figure 8.**
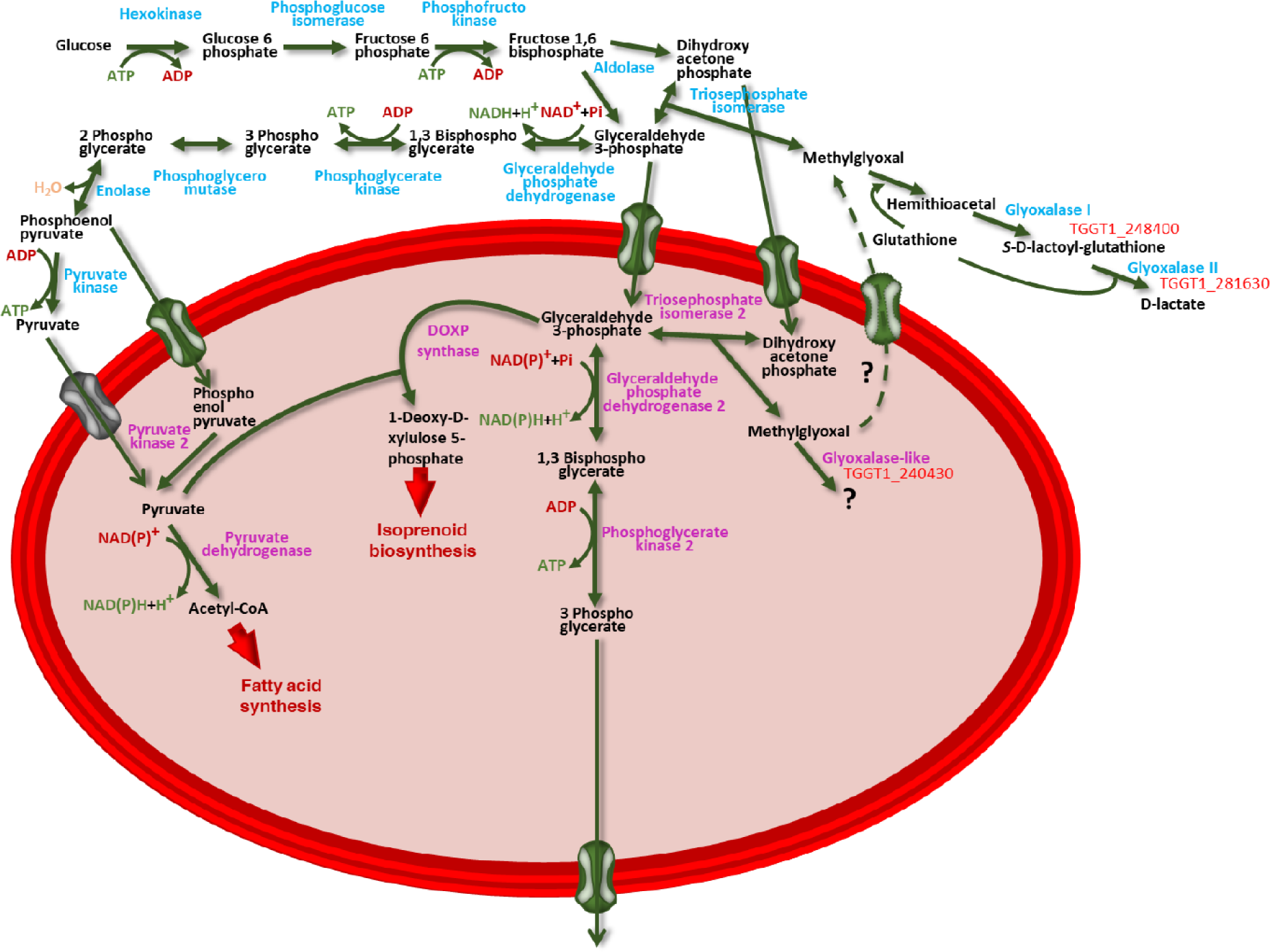
Shematic representation of the apicoplast-located metabolism of glycolytic intermediates. The cytosolic TgGlo2 (Toxodb.org entry TGGT1_281630) and the apicoplast-localized TgGloL (Toxodb.org entry TGGT1_240430) investigated in this study are mentioned.

Strikingly, both *P. falciparum* and *T. gondii* express a Glo1-domain containing protein in the apicoplast (PfGilp and TgGloL, respectively), with the addition of a Glo2-like protein also found in the organelle in *P. falciparum,* although the latter has no clear homologue in *T. gondii* (Fig. 1A). While this would argue for the presence of a local detoxification system in the apicoplast, previous studies in *P. falciparum* found no Glo1 activity associated with PfGilp (Akoachere *et al*., 2005) and lack of conservation of some potentially important amino acids in the active site both for PfGilp and TgGloL (Fig. 1D) would rather suggest that they may not function as Glo enzymes. It is also puzzling that these two Glo1 domain-containing proteins, while both expressed in the apicoplast of relatively close organisms, have very little homology between them and are quite phylogenetically divergent (Fig. 1B). This does not point to a conserved and ancient function that would be specifically associated with an apicoplast-dependent metabolism. Yet, while amino acids degeneracy may indicate loss of function during evolution, it may also reflect some lineage-specific specialization. Importantly, our data show unambiguously that TgGloL has an essential function for the fitness of *T. gondii* tachyzoites (Fig. 4), and that its depletion has a marked effect on the apicoplast (Fig. 6), for which it thus still performs an important role. The kinetics for the onset of the observed impact on the organelle, happening several days after TgGloL depletion, may be compatible with the progressive accumulation of a toxic compound. Yet, we could not detect any obvious accumulation of MG (Fig. 7A, B) or provide experimental evidence that the phenotype is due to a specific impairment in MG detoxification (Fig. 7).

TgGloL function at the apicoplast could then be completely unrelated to MG detoxification, although no specific domain or motif can be identified apart from the Glo1-like domains, even when using structure-based function annotation software. However, the βαβββ arrangement of Glo1-like domains present in TgGloL (Fig. 1C) is in fact a hallmark of the larger VOC family of metalloenzymes, that coordinate a divalent metal center that will usually bind to a ligand activate adjacent oxygen atoms (Armstrong, 2000; He and Moran, 2011). Beyond this common mechanistic attribute, these structurally-related enzymes are largely unrelated in terms of sequence, and are involved in very functionally diverse biological reactions: they include isomerases (like Glo1) and enzymes performic nucleophilic addition (like the Fosfomycin Resistance Protein), but also dioxygenases (like the extradiol dioxygenase, an oxidoreductase cleaving aromatic rings). TgGloL does not bear any particular homology to other VOC superfamily member besides Glo1 that would allow assigning a different function. We have also shown that, like PfGilp, TgGloL potentially lacks some metal/glutathione binding sites typically conserved in VOC enzymes (Fig. 1D), which raises questions about its ability to be metabolically active. However, it could have a mode of action similar to the Bleomycin Resistance Protein, which is also a member of the VOC superfamily that is neither an enzyme nor a metalloprotein, but for which the βαβββ fold creates a hydrophobic cavity that accommodates the antibiotic (Dumas *et al*., 1994). Thus, interestingly, there is a possibility that TgGloL has evolved to act as a sequestering protein for some metabolite (whether it is MG or not) that might build up in metabolically active apicoplasts.

There are still many outstanding questions as to how apicoplast-generated MG would be detoxified and this certainly deserves further investigation. The apicoplast is a validated drug target, and while most strategies aim at inhibiting the production of vital metabolites, inducing the accumulation of toxic intermediates that already lie within an endogenous metabolic pathway can be a very interesting alternative.

## Materials and methods

### Parasites and cells culture

Tachyzoites of the TATi ΔKu80 *T. gondii* strain (Sheiner *et al*., 2011) and derived transgenic parasites generated throughout this study were maintained by serial passage in monolayers of human foreskin fibroblast (HFF, American Type Culture Collection, CRL 1634) grown in Dulbecco’s modified Eagle medium (Gibco), supplemented with 5% decomplemented fetal bovine serum, 2-mM L-glutamine and a cocktail of antibiotics (penicillin-streptomycin) at 100 μg/ml.

### Bioinformatic analyses

Sequence alignment was performed using the CLUSTAL algorithm of the Geneious software suite v6.1.8 (http://www.geneious.com).

Phylogenetic analyses were performed with the MEGA software v11.0.13 package (https://www.megasoftware.net) (Tamura *et al*., 2021) as follows: whole sequences were aligned using the CLUSTAL method with a gap opening penalty of 10 and a gap extension penalty of 0.10, then the tree was built using the Maximum Likelihood method and Poisson correction model. Branches corresponding to partitions reproduced in less than 50% bootstrap replicates are collapsed. The analysis involved 11 amino acid sequences, with a total of 1119 positions in the final dataset.

Structural modelling was performed with AlphaFold (https://alphafold.ebi.ac.uk/) (Jumper *et al*., 2021) and structure visualization as well as structural comparisons were done with Swiss model (https://swissmodel.expasy.org/) (Waterhouse *et al*., 2018) and ChimeraX v1.8 (https://www.cgl.ucsf.edu/chimerax/) (Meng *et al*., 2023).

### Generation of the HA-tagged TgGloL cell line

A CRISPR-based strategy was used to tag the TgGloL protein (entry TGGT1_240430 in the www.toxodb.org database) by editing the native locus of the corresponding gene. Using the pLIC-HA_3_-CAT plasmid as a template, a PCR was performed with the KOD DNA polymerase (Novagen) to amplify a fragment with the tag and the resistance gene expression cassette with primers ML5311/ML5312 (all primers used in this work are listed in Table S1), which also carries 30 bp homology with the 3′ end of the corresponding genes. A specific single-guide RNA (sgRNA) was generated to introduce a double-stranded break at the 3′ of the *TgGloL* gene, using primers ML5317/ML5318, and the protospacer sequence was introduced in the Cas9-expressing pU6-Universal plasmid (Addgene, ref #52694) (Sidik *et al*., 2016). The TATi ΔKu80 cell line was transfected and transgenic parasites were selected with chloramphenicol and cloned by serial limiting dilution. Parasites with proper integration, named TgGloL-HA, were checked by PCR using primers ML5315/ML5489.

### Generation of the TgGloL conditional knock-down cell line

The conditional knock-down cell line for *TgGloL* was generated based on the Tet-Off system using the DHFR-TetO7Sag4 plasmid (Morlon-Guyot *et al*., 2014). A 1,504 bp fragment corresponding to the 5’ region of the gene, starting with the initiation codon, was amplified from genomic DNA by PCR using Q5 polymerase (New England Biolabs) with primers ML5313/ML5314 and cloned into the DHFR-TetO7Sag4 plasmid, downstream of the anhydrotetracycline (ATc)-inducible TetO7Sag4 promoter, yielding the DHFR-TetO7Sag4-TgGloL plasmid. The plasmid was then linearized and transfected into the TgGloL-HA cell line. Transfected parasites were selected with pyrimethamine and cloned by serial limiting dilution, yielding the cKD TgGloL-HA cell line, in which proper integration was verified by PCR using primers ML1041/ML5531.

### Generation of the EcGlo3-expressing TgGloL conditional knock-down cell line

A 1,206 bp cassette containing at its 5’ 324 bp coding for a 108 amino acid-long N-terminal sequence sufficient for targeting *T. gondii* ferredoxin NADP^+^ reductase (www.toxodb.org entry TGGT1_298990) to the apicoplast lumen (Harb *et al*., 2004), fused to the coding sequence for *E. coli* Glo3 (or protein/nucleic acid deglycase 1; www.uniprot.org entry P31658), fused to a 3’ sequence coding for a TY tag (Bastin *et al*., 1996), was synthesized by GenScript (www.genscript.com). The cassette was cloned into the pUPRT-TUB-Ty vector (Sheiner *et al*., 2011). This plasmid was subsequently linearized and cotransfected together with a plasmid expressing Cas9 and a guide RNA specific of the *uracil phosphoribosyl transferase* (*UPRT*) locus under the control of a U6 promoter (Nguyen *et al*., 2018) in the cKD TgGloL-HA cell line. Transgenic parasites were selected with 5-fluorodeoxyuridine, cloned by serial limiting dilution, and checked by PCR with primers ML6396/ML6397.

### Generation of the HA-tagged TgGlo2 cell line

A CRISPR-based strategy was used to tag the TgGlo2 protein (entry TGGT1_281630 in the www.toxodb.org database) by editing the native locus of the corresponding gene. Using the pLIC-HA_3_-CAT plasmid as a template, a PCR was performed with the KOD DNA polymerase (Novagen) to amplify a fragment with the tag and the resistance gene expression cassette with primers ML5907/ML5908 (all primers used in this work are listed in Table S1), which also carries 30 bp homology with the 3′ end of the corresponding genes. A specific single-guide RNA (sgRNA) was generated to introduce a double-stranded break at the 3′ of the *TgGlo2* gene, using primers ML5905/ML5906, and the protospacer sequence was introduced in the Cas9-expressing pU6-Universal plasmid (Addgene, ref #52694) (Sidik *et al*., 2016). The TATi ΔKu80 cell line was transfected and transgenic parasites were selected with chloramphenicol and cloned by serial limiting dilution. Parasites with proper integration, named TgGlo2-HA, were checked by PCR using primers ML5904/ML1476.

### Immunoblot analysis

Protein extracts from 10^7^ freshly egressed tachyzoites were prepared in Laemmli sample buffer, separated by SDS-PAGE on a 10% acrylamide gel and transferred onto nitrocellulose membrane using a Mini-Transblot system (BioRad) according to the manufacturer’s instructions. Membranes were blocked in Tris-Buffered Saline with 0.1% v/v Tween-20 and 5% w/v nonfat dry milk and were subsequently incubated with primary and secondary antibodies in the same solution. Primary antibodies used were: rat monoclonal anti-HA tag (clone 3F10, Roche) which was used at 1/500, mouse anti-actin (Herm-Gotz, 2002) (hybridoma supernatant used at 1/25), rabbit anti-PDH E2 (Renaud *et al*., 2022) which was used at 1/500, mouse anti-TY tag (Bastin *et al*., 1996) used at 1/500, mouse monoclonal anti-AGE antibody (clone 3D11, Merck) and mouse monoclonal Anti-methylglyoxal (clone 9F11, Merck), which were used at 1/200.

### Immunofluorescence microscopy

For immunofluorescence assays (IFA), intracellular tachyzoites grown on coverslips containing HFF monolayers, were either fixed for 20 min with 4% (w/v) paraformaldehyde in PBS and permeabilized for 10 min with 0.3% Triton X-100 in PBS or fixed for 5 min in cold methanol (for SAG labeling). Slides/coverslips were subsequently blocked with 1% (w/v) BSA in PBS. Primary antibodies used (at 1/1,000, unless specified) were rat anti-HA tag (clone 3F10, Roche), rabbit anti-Cpn60 (Agrawal *et al*., 2009), rabbit anti-IMC3 (Anderson-White *et al*., 2011), mouse anti-SAG1 (Couvreur *et al*., 1988), and mouse anti-SAG3 (Cesbron-Delauw *et al*., 1994), mouse anti-F1β ATPase (gift of P. Bradley), mouse anti-MIC3 (clone T4 2F3) (Garcia-Réguet *et al*., 2000), mouse anti-TY tag (Bastin *et al*., 1996), mouse anti-AGE/methylgyoxal antibodies (clones 3D11 and 9F11, respectively, Merck) used at 1/200. Staining of DNA was performed on fixed cells by incubating them for 5 min in a 1 μg/ml 4,6-diamidino-2-phenylindole (DAPI) solution. All images were acquired at the Montpellier RIO imaging facility with a Zeiss AXIO Imager Z2 epifluorescence microscope driven by the ZEN software v2.3 (Zeiss). Z-stack acquisition and maximal intensity projection was performed to quantify apicoplast loss. Adjustments for brightness and contrast were applied uniformly on the entire image.

### Electron Microscopy

Parasites were pretreated for three days with ATc, and then used to infect HFF monolayers and grown for an extra 24 hours in ATc. Untreated parasites were used as a control for normal morphology. They were fixed with 2.5% glutaraldehyde in cacodylate buffer 0.1 M pH7.4. Coverslips were then processed using a Pelco Biowave pro+ (Ted Pella). Briefly, samples were postfixed in 1% OsO_4_ and 2% uranyl acetate, dehydrated in acetonitrile series and embedded in Epon 118 using the following parameters: Glutaraldehyde (150 W ON/OFF/ON 1-min cycles); two buffer washes (40 s 150 W); OsO_4_ (150 W ON/OFF/ON/OFF/ON 1-min cycles); two water washes (40 s 150 W); uranyl acetate (100 W ON/OFF/ON 1-min cycles); dehydration (40 s 150 W); resin infiltration (350 W 3-min cycles). Fixation and infiltration steps were performed under vacuum. Polymerization was performed at 60°C for 48 hr. Ultrathin sections at 70 nM were cut with a Leica UC7 ultramicrotome, counterstained with uranyl acetate and lead citrate and observed in a Jeol 1400+ transmission electron microscope from the MEA Montpellier Electron Microscopy Platform. All chemicals were from Electron Microscopy Sciences, and solvents were from Sigma.

### Plaque assay

Confluent monolayers of HFFs were infected with freshly egressed parasites, which were left to grow for 7 days in the absence or presence of ATc (unless stated). They were then fixed with 4% v/v paraformaldehyde (PFA) and plaques were revealed by staining with a 0.1% crystal violet solution (V5265, Sigma-Aldrich). Plaques were imaged with an Olympus MVX10 stereomicroscope and their area was measured using the Zen 2.3 software (Zeiss) with the “contour” tool.

### Statistical analyses

All statistical analyses were performed with the Prism 8 software (Graphpad) using two-tailed Student’s *t*-test comparisons.

## Acknowledgements

We thank MJ. Gubbels, P. Bradley, D. Soldati-Favre V. Carruthers and S. Lourido for providing plasmids and antibodies. We also thank M. Gonzalez-Durany and Yann Bordat for technical help in the early stages of the project. We acknowledge the Toxodb/Veupathdb databases, their curators and their contributors for invaluable source of data. This work was supported by the Agence Nationale de la Recherche (grants ANR-19-CE15-0023 and ANR-22-CE20-0026).

## Figures

**Table S1. Primers used in this study.**

## Notes

### Competing Interest Statement

The authors have declared no competing interest.

